# Regulation of Phosphatidylinositol-(4,5)-bisphosphate and Active-Rho1p Levels and Distribution is Crucial for Correct Spatio-temporal Cytokinesis and Echinocandin Responses in *Candida albicans*

**DOI:** 10.1101/2024.12.18.629289

**Authors:** Hassan Badrane, M. Hong Nguyen, Cornelius J. Clancy

## Abstract

*Candida* species cause severe infections like invasive candidiasis, which annually affect 1.5 million people worldwide and cause close to 1 million deaths. *Candida albicans* is the predominant cause of candidiasis. We previously showed that EH domain-containing protein Irs4p binds 5-phosphatase enzyme Inp51p to regulate plasma membrane levels of phosphatidylinositol-(4,5)-bisphosphate (PI(4,5)P_2_) in *C. albicans*. Indeed, deletion of *IRS4* or *INP51* led to elevated levels of PI(4,5)P_2_ and presence of abnormal intracellular membranous PI(4,5)P_2_ patches. We demonstrated an interplay between PI(4,5)P_2_ and septins to regulate the PKC-Mkc1 cell wall integrity pathway, echinocandin and cell wall stress responses, and virulence during candidiasis. In the current investigation, we used fluorescent protein tagging and live cell imaging to follow the nascency of PI(4,5)P_2_ patches. We show that these abnormal patches tightly correlate with cytokinesis, as they predominantly arise close to the site and time of cell division. We further demonstrate these patches colocalize PI(4,5)P_2_ with actomyosin ring components Act1p and Myo1p, which form its core, and active Rho1p, a small GTPase that plays a regulatory role. Additionally, activation of Rho1p was altered in *irs4* and *inp51* mutants compared to wild-type strain, with over-activation or down-activation during early exponential or stationary phase, respectively. Wild-type cells exposed to 4xMIC of the echinocandin caspofungin show abnormal PI(4,5)P_2_ patches colocalizing the same cytokinesis components as above, except that they were transient. Taken together, our results support a model in which PI(4,5)P_2_ plays a pivotal role, along with Rho1p, in the correct execution of cytokinesis and response to caspofungin.

## INTRODUCTION

*Candida* infections are caused by diverse commensal, opportunistic budding yeast species belonging to distinct families in the *Saccharomycetales* order (1). These infections manifest different clinical forms ranging from non-life threatening superficial mucocutaneous infections like oropharyngeal and vulvovaginal candidiasis, or superficial skin infections, to the potentially lethal deep-seated and disseminated invasive candidiasis (IC). Mortality rates from IC can reach 20%-40% for immunocompromised individuals and hospitalized patients with serious underlying diseases, and they pose a significant economic burden to healthcare systems (2). *Candida albicans* is the leading cause of IC. Echinocandins are the agents of choice against most infections by *C. albicans* and other *Candida* spp. These drugs exert fungicidal activity on *Candida* by damaging the cell wall through inhibition of 1,3-β-D-glucan synthase, a critical enzyme in cell wall biogenesis.

The *Candida* cell wall and adjoining plasma membrane (PM) are crucial to pathogenesis of IC and echinocandin responsiveness (3). Phosphatidylinositol-(4,5)-bisphosphate (PI(4,5)P_2_) localizes to the PM’s cytoplasmic layer, where it anchors via its fatty acid chain and glycerol backbone (4). This leaves its inositol headgroup available to interact conformationally with PH domain proteins (5) or electrostatically with basic residues of arginine and lysine (6). In previous work, we found that exposure of wild-type *C. albicans* cells to the echinocandin caspofungin led to an elevation of PI(4,5)P_2_ levels in a rapid and dose-dependent fashion, and a dynamic mislocalization of both PI(4,5)P_2_ and septins within aberrant PM invaginations (7). Many *C. albicans* cells exposed to caspofungin also demonstrated abnormally wide bud-necks (8). In other studies, we showed that the *C. albicans* EH domain-containing protein Irs4p binds the 5-phosphatase enzyme Inp51p to regulate intracellular levels of PI(4,5)P_2_. *C. albicans* mutant strains with deletion of genes encoding either Irs4p or Inp51p over-activated the PKC-Mkc1 cell wall integrity pathway upon caspofungin exposure, and they were hyper-susceptible to the drug and attenuated for virulence in a hematogenous disseminated mouse model of IC (9, 10). Mutant strains had elevated levels of PI(4,5)P_2_ and PI(4,5)P_2_-containing PM invaginations that resembled those of wild-type *C. albicans* exposed to caspofungin. Abnormal invaginations also encompassed septins (Sep7p and Cdc10p), chitin, GPI-anchored proteins (Rbt5p), and cell wall material (7, 9, 10). Taken together, our previous data indicate that PI(4,5)P_2_-septin regulation is crucial in echinocandin and cell wall integrity responses, and in *C. albicans* virulence.

While many research investigations established the involvement of PI(4,5)P_2_ or it’s degradation, in several functions like cytoskeleton remodeling, endocytosis, exocytosis, ion channel regulation, and signaling pathways ((4, 11) for review), only a handful of studies provided evidence hinting to the possible role of PI(4,5)P_2_ in cytokinesis (12). In the current study, we provide the first strong evidence implicating PI(4,5)P_2_ in cytokinesis in *C. albicans*. And we further show its importance for echinocandin responses. Using live cell imaging, we first looked at the spatio-temporal connection between the appearance of PI(4,5)P_2_ patches and cytokinesis in *irs4* and *inp51* mutant cells. Then, we examined co-localization of PI(4,5)P_2_ in the mutants with major components of the actinomyosin ring (Act1p and Myo1p) and with the GTP-binding protein Rho1p, which is an upstream activator of the PKC-Mkc1 cell wall integrity pathway, a regulator of the actin cytoskeleton during cytokinesis, and a regulatory subunit of 1,3-β-D-glucan synthase. Finally, we determined if abnormalities in mutant cells were also evident in wild-type *C. albicans* cells exposed to caspofungin.

## MATERIALS & METHODS

### Strains, Growth Conditions, PCR, and subcloning

*C. albicans* strains used in this study are listed in Table 1. Strains were grown in Yeast-Peptone-Dextrose (YPD) or Synthetic Defined (SD) Media at 30°C. All oligonucleotides used in this study are available upon request. PCR was performed using high-fidelity polymerases, including Agilent’s pfuUltra II Fusion HS DNA polymerase (Santa Clara, CA, USA) or, for long PCR fragments (5kb-12kb), Takara’s PrimeSTAR Max DNA polymerase (Mountain View, CA, USA). For on-colony PCR screening, we used 5PRIME HotMaster Taq DNA Polymerase (Quantabio, Beverly, MA, USA). PCR conditions were per manufacturer recommendations, overall annealing temperature was 54°C-58°C, and the elongation time was 15s-30s/kb. To perform chimeric protein fusion, PCR bands were purified with Nucleospin Gel and PCR Clean-up Kit (Takara BIO USA), and ligated with Quick Ligase Kit (New England Biolabs, Ipswich, MA, USA). The ligated PCR fragments ends should have corresponding oligos with a 5’ phosphorylation. Nested PCR was performed to amplify the fused ORFs, using primers carrying appropriate restriction enzyme sites for subcloning into pSFS2A plasmid (13). We previously integrated into this plasmid: CaADH1 promoter and terminator to control the expression of the fused ORFs, F1 and F2 DNA fragments for integration by recombination into a noncoding genomic region at coordinates between 625,000 and 627,000 of chromosome 1 upstream of RP10 locus, as well as either CaGFP or CaRFP for fluorescent tagging (7, 8). NEB 10-beta chemically competent *E. coli* were used for plasmid transformation (New England Biolabs, Ipswich, MA, USA), which was carried out following the manufacturer’s heat-shock protocol. This *E. coli* strain was particularly efficient in transforming large DNA (10kb – 14kb).

**TABLE 1.**
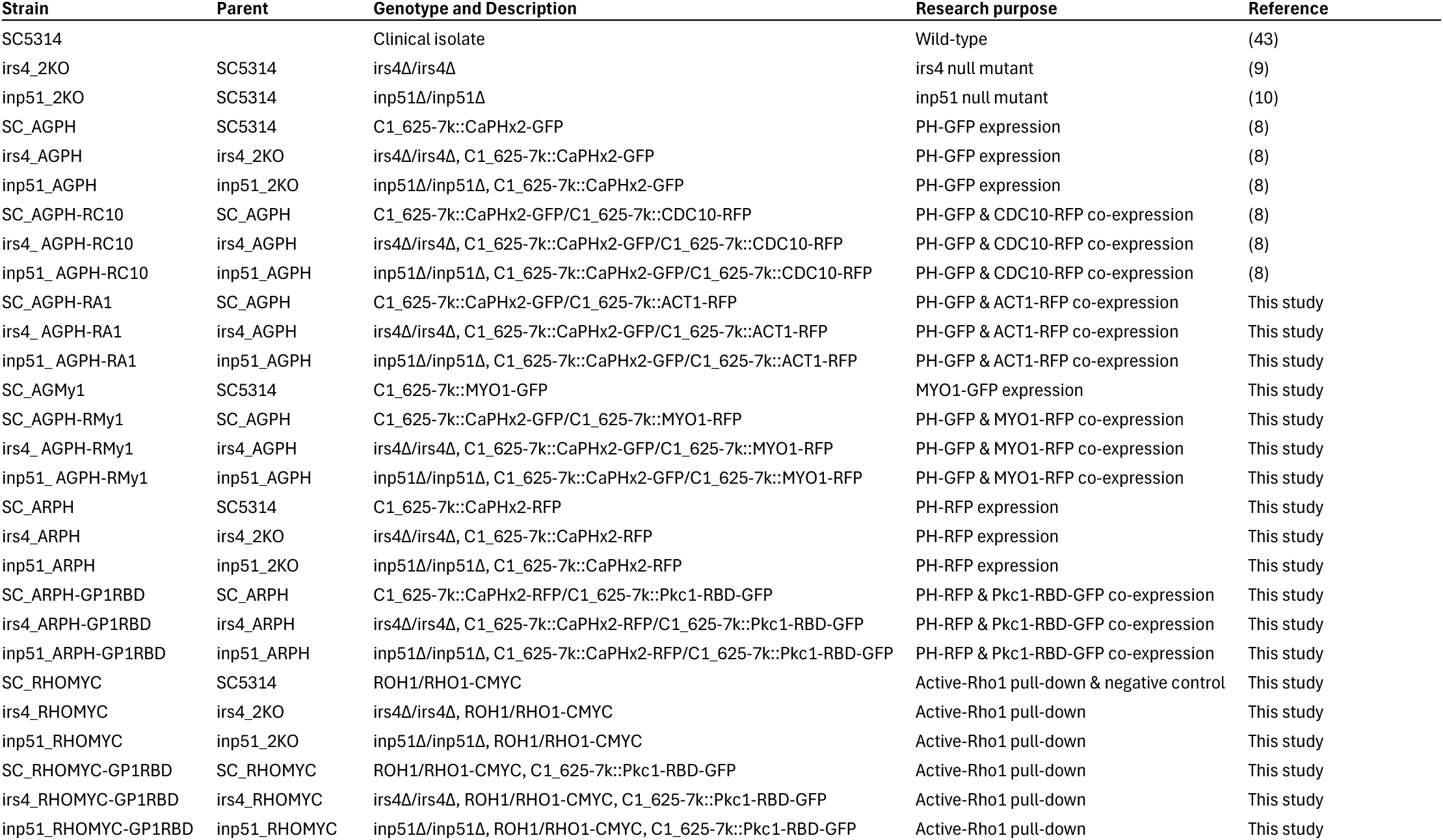
List of strains used in this study.

### Transformation

DNA for transformation was prepared by digestion of plasmid followed by gel purification of the band of interest, or by PCR amplifying the whole cassette (10k – 14kb) using Takara’s PrimeSTAR Max DNA polymerase with M13 and T7 primers, then purifying the PCR band. Transformation of *C. albicans* was done using an electroporation protocol described previously (13). Transformants were selected on YPD plates supplemented with 200 µg/ml of Nourseothricin (13). Positive transformants were screened by microscopy and PCR.

### Live cell imaging

*C. albicans* cells were grown overnight in YPD medium at 30°C. On the next day, cells were subcultured in fresh YPD at 30°C for 3-4 hours and 150 μl of culture were deposited on a 35-mm glass-bottom dish (Matek, Ashland, MA). The dishes were pretreated with 10 μg/cm^2^ of Cell-Tak adhesive (BD Biosciences, Bedford, MA) per manufacturer instructions, to adsorb a thin layer of polyphenolic proteins on which cells were immobilized for microscopy. Dishes with the culture were incubated at 30°C for 1 hour, then the medium was removed and cells were washed twice with sterile DDW and once with fresh YPD to remove unattached cells. Finally, 150 μl of fresh YPD medium (or YPD supplemented with appropriate amount of caspofungin) were added and the dishes were placed in the heated stage of the microscope. Microscopy was performed at the University of Pittsburgh Center for Biologic Imaging, using established protocols (7, 8). We used a Nikon A1 confocal microscope for live cell imaging and acquired data with NIS Elements software (Nikon, Minato, Tokyo, Japan).

### Pulldown assay for active Rho1

Similarly to the subcloning method described above, *RHO1* was fused to a 13xcMyc tag, and a construct was generated to replace the native gene with the tagged version using the SAT flipper tool (13). After transforming cells with this cassette, transformants were screened with PCR to ensure correct integration. Further, we used Western Blot as described before (8) with anti-cMyc monoclonal antibody 9E10 (Thermo Fisher Scientific, Waltham, MA, USA) to confirm expression and correct size of the tagged protein. On the other side, Rho1 binding domain from *C. albicans PKC1* (PKC-RBD; (14)) was amplified and fused to CaGFP, and transformed for coexpression with Rho1p-13xcMyc. Cells were grown and harvested after overnight culture or after an additional 4 hours subculture. Proteins were extracted by cell disruption using a Mini-Beadbeater-8 (Biospec, Bartlesville, OK, USA), in NP-40 lysis buffer supplemented with protease inhibitors, Protease Inhibitor Cocktail and PMSF (Sigma-Aldrich, Saint-Louis, MO, USA). Active Rho1-13xMyc, interacting with PKC-RBD-CaGFP, was pulled down using the GFP-Trap Agarose kit (Chromotek, Planegg, Germany). Finally, Western Blot with 9E10 anti-cMyc monoclonal antibody was used to detect active Rho1p. A strain expressing Rho1-13xMyc alone, without the PKC-RBD-CaGFP, was used with the GFP-Trap Agarose kit as a pulldown negative control.

## RESULTS

### PI(4,5)P_2_ patches correlate with cytokinesis in space and time

Mutant *C. albicans inp51* and *irs4* cells expressing the pleckstrin homology (PH) domain fused to CaGFP were used for live cell imaging with confocal microscopy. We have previously shown that this PH domain, which is encoded by the human Phospholipase C-δ1 (PLC), binds specifically *in vivo* to PI(4,5)P_2_ in *C. albicans* (7, 15, 16). Generally, about 50% of cells showed the PI(4,5)P_2_ patches. We followed these abnormal PI(4,5)P_2_ patches and scored their appearance with regards to proximity in space to the site of cell division or concurrence during cytokinesis. There was a tight association between abnormal PI(4,5)P_2_ patches and cytokinesis, as 83% and 84% of them appeared during and in proximity to cytokinesis, respectively (Table 2). Only twelve percent of them were not associated with either cytokinesis sites or timing. Representative examples of these events are shown in Fig. 1, and in time lapse videos (Supplemental Video 1a and 1b). These observations strongly link these abnormal PI(4,5)P_2_ patches with cytokinesis in time and space.

**FIG 1.**
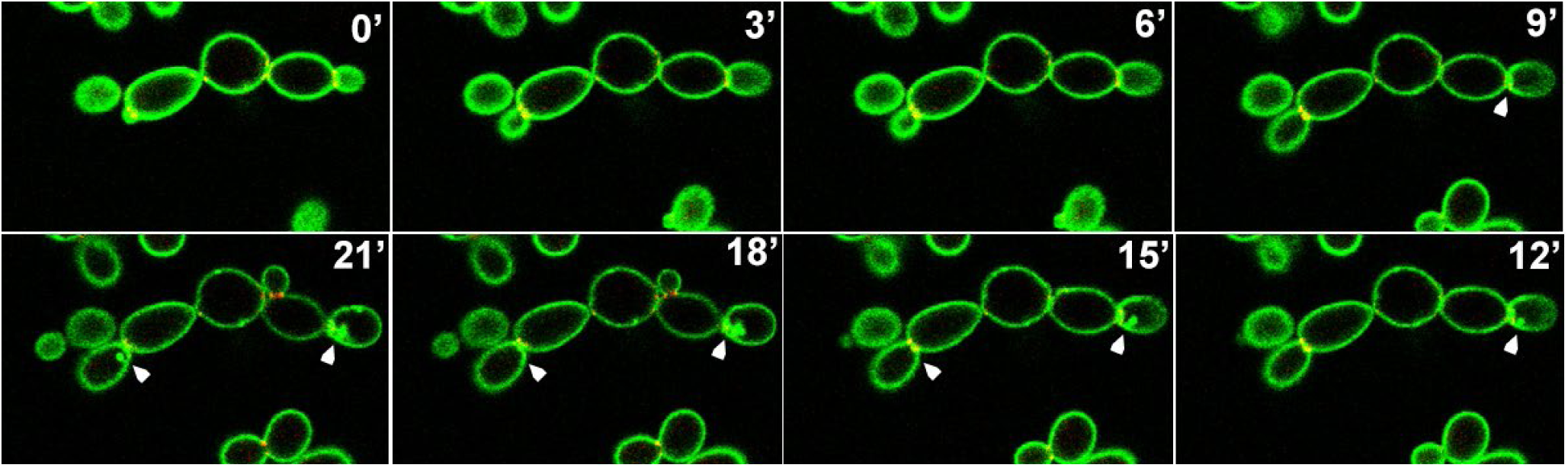
The abnormal PI(4,5)P_2_ patches are linked to cytokinesis in space and time. Three minutes interval time lapse of dividing cells from *inp51* mutant (similar results with *irs4*) expressing CaPHx2-CaGFP and CDC10-CaRFP, showing dynamics of the appearance of the abnormal PI(4,5)P_2_ patches, which are formed close to cytokinesis in space and time. White arrows are depicting two division sites in which the abnormal PI(4,5)P_2_ patches appear right after the end and close to the site of cytokinesis (See Supplemental Video 1a and 1b).

**TABLE 2.** Scoring of PI(4,5)P_2_ patches with regards to cytokinesis spatio-temporal dynamics. Shown is each case are number of occurences (percentage)

|  | **Cytokinesis Site** | **Elsewhere** | **TOTAL** |
| --- | --- | --- | --- |
| **During Cytokinesis** | 87 (78%) | 6 (5%) | 93 (83%) |
| **Not during Cytokinesis** | 7 (6%) | 12 (11%) | 19 (17%) |
| **TOTAL** | 94 (84%) | 18 (16%) | 112 (100%) |

### Act1p and Myo1p colocalize with the abnormal PI(4,5)P_2_ patches

Actin and type II myosin heavy chain are encoded by *ACT1* and *MYO1*, respectively, and are highly conserved across animal and yeast species. They are the main components of actomyosin, which forms the contractile ring that functions and is essential during cytokinesis (Reviewed in (17)). We transformed mutant cells expressing CaPHx2-CaGFP with a cassette to constitutively co-express CaACT1-CaRFP. Using a similar approach, we previously showed that septins (CaSep7p and CaCDC10p) co-localized with abnormal PI(4,5)P_2_ patches in mutant strains. In wild-type cells, Act1p was found localized to cortical actin patches, as expected. In mutants, however, Act1p was mislocalized and did colocalize with the abnormal PI(4,5)P_2_ patches (Fig. 2, Supplemental Video 2a and 2b). Likewise, we co-expressed CaMYO1-CaRFP to localize myosin. In Live cell confocal imaging of wild-type cells, Myo1p localized initially to the bud neck, then translocated from mother cell to the apical tip of the daughter cell during S-phase. At the end of anaphase and just before completion of cell division, Myo1p migrated to the site of cell division as cytokinesis is completed (See Supplemental Video 3a and 3b). In *irs4* or *inp51* mutant cells during cytokinesis, Myo1p is mislocalized and co-localized with abnormal PI(4,5)P_2_ patches as the patches emerge (Fig. 3, Supplemental Video 3c). These data show the main components of actomyosin, an essential element of cytokinesis machinery, to be mislocalized with PI(4,5)P_2_ patches.

**FIG 2.**
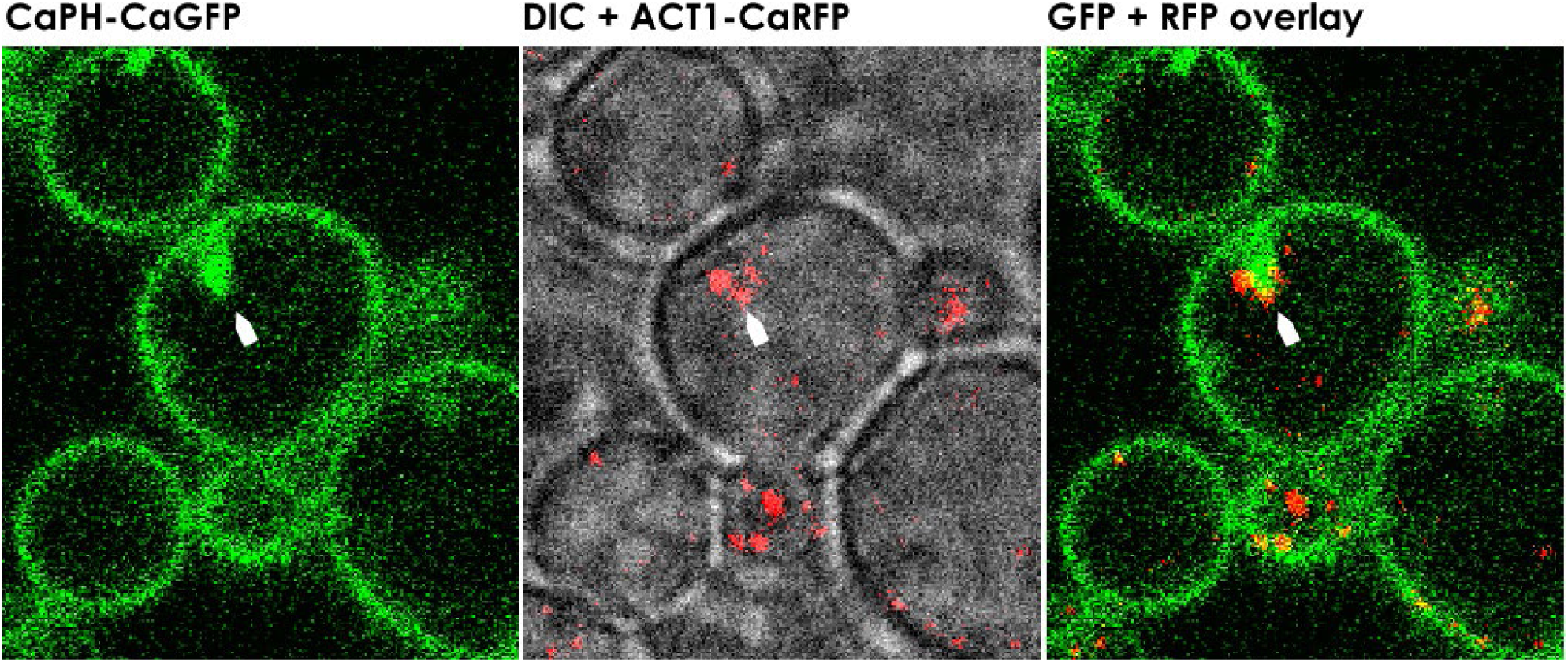
Act1p colocalizes with the abnormal PI(4,5)P_2_ patches. Confocal microscopy of *irs4* (or *inp51*) mutant expressing CaPHx2-CaGFP and ACT1-CaRFP. Left panel shows GFP signal, center panel shows a merging of DIC and RFP signal, and right panel shows merging of GFP and RFP signals. White arrows depict co-mislocalization of PI(4,5)P_2_ and Act1p in the abnormal PI(4,5)P_2_ patches (See Supplemental Video 2a and 2b).

**FIG 3.**
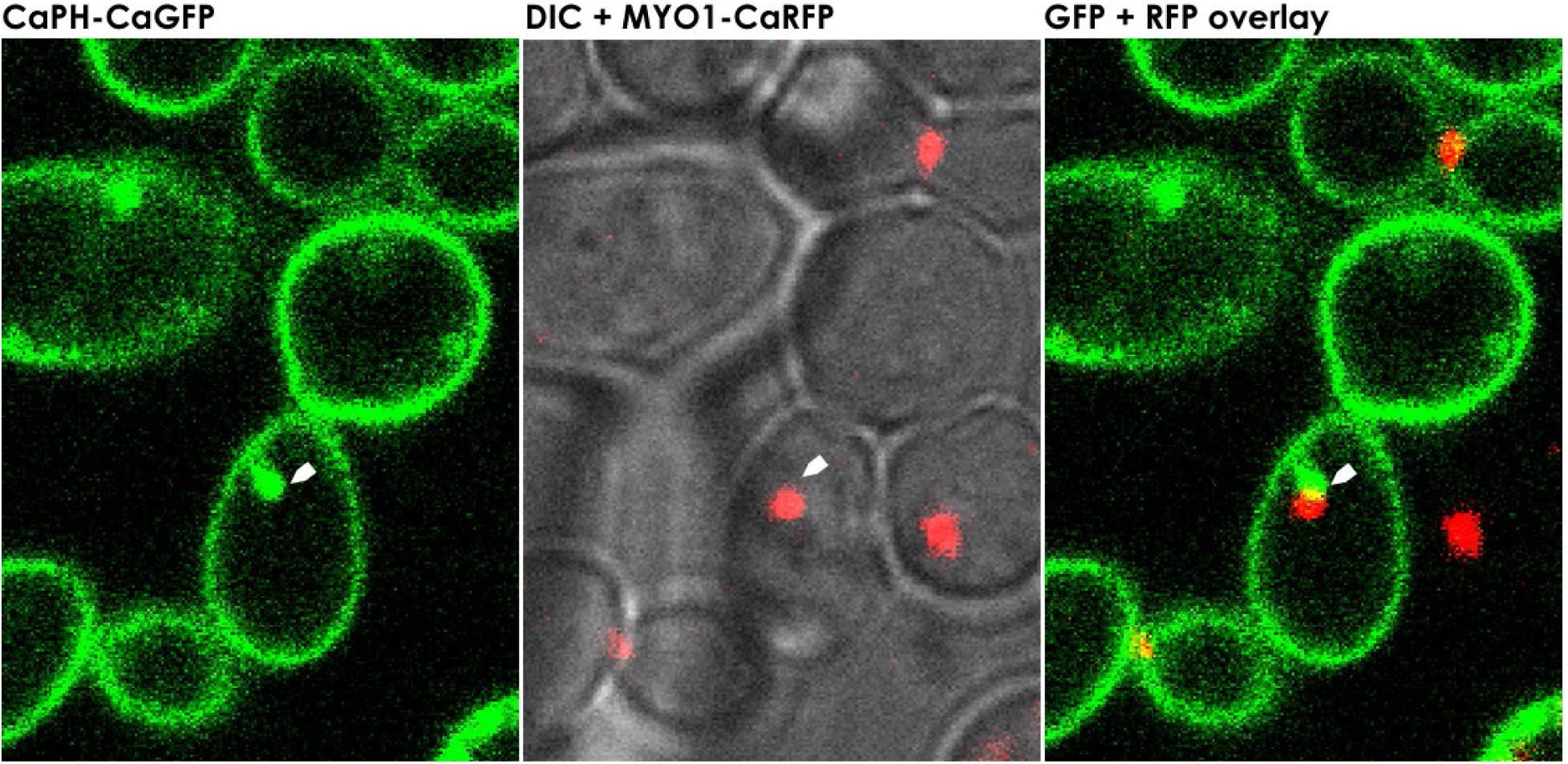
Myo1p colocalize with the abnormal PI(4,5)P_2_ patches. Confocal microscopy of *irs4* (or *inp51*) mutant expressing CaPHx2-CaGFP and CaMYO1-CaRFP. Left panel shows GFP signal, center panel shows a merging of DIC and RFP signal, and right panel shows merging of GFP and RFP signals. White arrows depict co-mislocalization of PI(4,5)P_2_ and Myo1p in the abnormal patches (See Supplemental Video 3c).

### Active-Rho1p is mislocalized to the abnormal PI(4,5)P_2_ patches, with altered activation in *irs4* or *inp51* mutant cells

In baker’s yeast, the small GTPase Rho1p, an essential protein, has been implicated in cell polarization, actin organization and cell wall synthesis (18–20). One of the effectors of Rho1p is Pkc1, a protein kinase that signals to the Mpk1 MAP kinase cascade to control actin cytoskeleton organization and cell wall biosynthesis genes (21–24). Pkc1p was shown to specifically bind active GTP-bound Rho1p via the Rho1 binding domain (RBD) (21). As a reporter for active Rho1, we fused PCR-amplified *C. albicans* Pkc1p-RBD (aa 371 to 636) to CaGFP (14) and transformed the cassette into wild-type cells. Live cell imaging of these transformed wild-type showed a GFP signal representing active Rho1p in the cell periphery, which intensified at the emerging bud (Supplemental Video 4a). Later, active Rho1p distribution constituted a gradient along the periphery of the daughter cell with increasing intensity toward the bud tip. This gradient faded progressively to disappear toward the end of the anaphase, where active Rho1p was evenly distributed along the cell periphery (Supplemental Video 4a). During cytokinesis, it was highly localized to the site of cell division where a new septum was formed. Then, active Rho1p decreased to normal levels at the end of cell division. Likewise, we observed the same distribution during hyphal growth (Supplemental Video 4b), as was also shown in previous studies (14).

Similarly, the Pkc1p-RBD-CaGFP cassette was transformed in wild-type or mutants backgrounds for coexpression either with Rho1p-13xcMyc, for pull-down assay, or with CaPHx2-CaRFP for subcellular co-localization using fluorescent imaging. Live cell confocal imaging of either *irs4* or *inp51* mutants co-expressing CaPHx2-CaRFP and PKC-RBD-CaGFP showed that in addition to the wild-type distribution of active Rho1p, there was an abnormal distribution that colocalized with aberrant PI(4,5)P_2_ patches (Fig. 4). The advent of this mislocalization happened mostly during the end of cytokinesis (Supplemental Video 5). In addition to abnormalities in the distribution of active Rho1p, we quantitatively investigated active Rho1p in pull-down experiments followed by western blot on cells co-expressing RHO1-13xcMyc and PKC-RBD-CaGFP. But firstly, we used a strain that only expressed RHO1-13xcMyc as a negative control. As expected, after pull-down with GFP-trap and western blot with α-Myc no band could be detected (data not shown). *Irs4* and *inp51* co-expressing RHO1-13xcMyc and PKC-RBD-CaGFP showed an under-activation of Rho1p compared to wild-type during stationary phase (Fig. 5, top panel). On the other hand, the opposite was observed during early exponential phase, as an overactivation of Rho1p was found in the mutants compared to wild-type (Fig. 5, bottom panel). Overall, these findings imply that a disruption in localization and levels of PI(4,5)P_2_ in mutants is accompanied by a disturbance in Rho1p regulation and localization of its active form.

**FIG 4.**
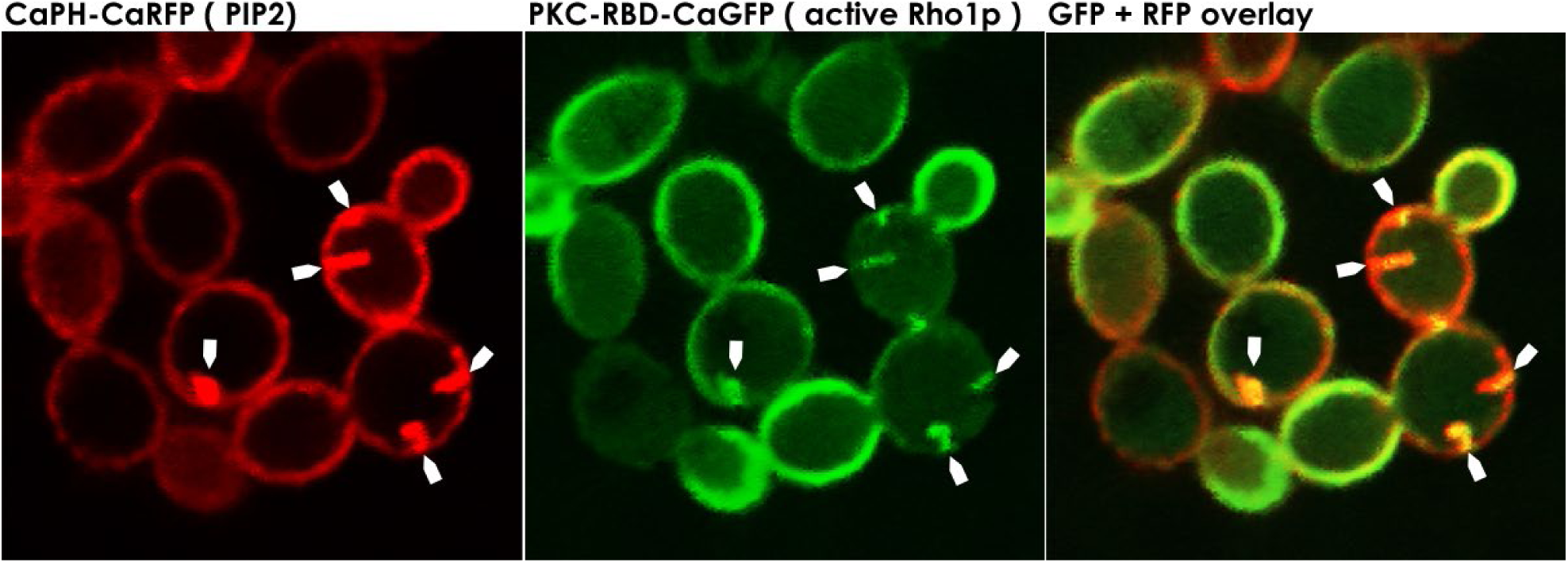
Active Rho1p mislocalizes with the abnormal PI(4,5)P_2_ patches. Confocal microscopy *inp51* (or *irs4*) mutant expressing CaPHx2-CaRFP and PKC-RBD-CaGFP. Left panel shows RFP signal representing distribution of PI(4,5)P_2_ in the PM, center panel shows GFP signal representing distribution of active-Rho1p, and right panel shows merging of GFP and RFP signals, where yellow signal represents colocalization of PI(4,5)P_2_ and active Rho1p in the abnormal patches (depicted by white arrows, See Supplemental Video 5).

**FIG 5.**
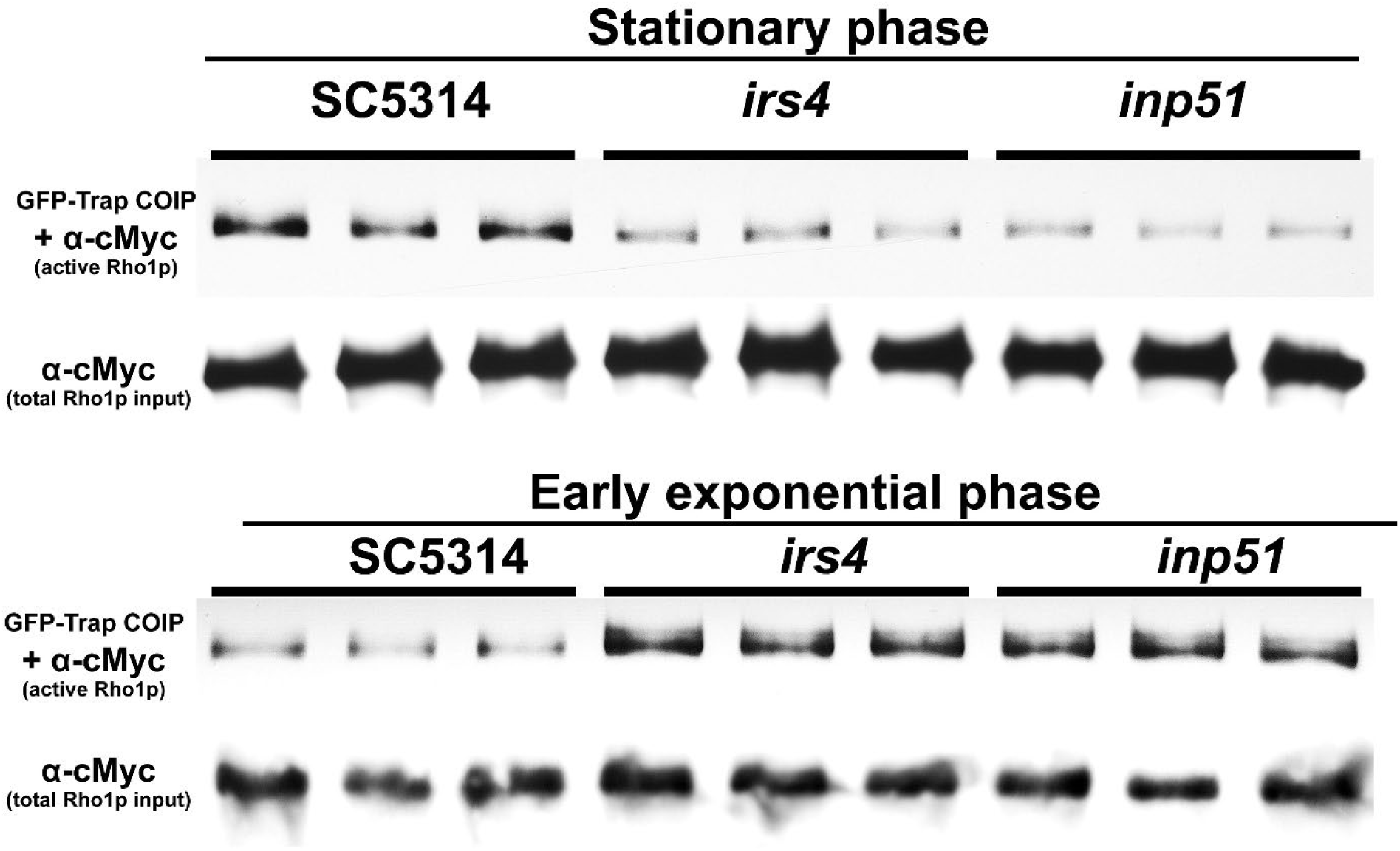
Mutants display an altered profile of Rho1p activation. Cells were grown overnight (stationary phase) or with an additional 4 hours subculture (early exponential phase). After harvesting, cells were disrupted, and cell lysates with equal total protein content (200 µg) were incubated with anti-GFP (GFP-Trap, Chromotek) to pull-down active Rho1p which binds PKC-RBD-CaGFP. A Western Blot using α-Myc was performed with pull-down eluate to detect active-Rho1p, or with 2 µg of total protein lysate to detect total Rho1p. Top panel shows a significant underactivation of Rho1p during stationary phase, while bottom panel shows a significant overactivation of Rho1p during early exponential phase.

### Similar defects related to PI(4,5)P_2_ are manifested during exposure of *Candida* wild-type cells to caspofungin

Our earlier study showed that wild-type *C. albicans* cells exposed to different concentrations of caspofungin rapidly mislocalized PI(4,5)P_2_ in a highly dynamic fashion and in a dose-dependent manner that correlated with fungicidal activity (7, 8). Caspfoungin exposure not only disturbed PI(4,5)P_2_ distribution but also augmented PI(4,5)P_2_ levels over a 3-hour exposure experiment. Our earlier study showed some abnormal septation events, and mislocalization of septins (Cdc10p and Sep7p) with PI(4,5)P_2_ (8). In this study, we exposed wild-type cells co-expressing CaPHx2-CaGFP and ACT1-CaRFP to 4xMIC Caspfoungin. Fig. 6 shows that actin was recruited to a site of the PM where it seemed to be enriching with PI(4,5)P_2_. Further, actin seemingly pulled the PM with PI(4,5)P_2_ inside the cell but later appeared to dissociate from the PM and PI(4,5)P_2_ (Fig. 6, Supplemental Video 6). Likewise, wild-type cells co-expressing CaPHx2-CaGFP and MYO1-CaRFP mislocalized Myo1p along with PI(4,5)P_2_ (Fig. 7, 8, Supplemental Video 7 and 8). Fig. 7 shows that Myo1p mislocalized with PI(4,5)P_2_ starting at 9’ right at the site of cell division and after completion of cytokinesis (Supplemental Video 7). The mislocalization started fading at 30’ when Myo1p dissociated from PI(4,5)P_2_. Fig. 8 shows another type of defect caused by exposure to caspofungin, as Myo1p localization was normal during cell division from 5’ to 80’ (indicated with white arrow), while PI(4,5)P_2_ showed multiple dynamic mislocalizations. During the following cell division at 95’, the daughter cell from previous cell division started a bud at a site (indicated by white arrow) but Myo1 is localized to a different site where another bud is started at 110’ (white arrowhead). The initial bud seemed to stall, but the second one continued the cell division with intermittent PI(4,5)P_2_ mislocalizations (Supplemental Video 8). The figure and supplemental video also show a wider mother-daughter bud neck, a cellular abnormality we previously observed with exposure to caspofungin (8). Finally, Fig. 9 shows dynamic mislocalization of both active-Rho1p and PI(4,5)P_2_, which together co-mislocalized to two different cellular localizations during a 6 min interval. All these data show strikingly similar mislocalizations of key components of cytokinesis after deletion of either *INP51* or *IRS4*, or after exposure to caspofungin. However, these phenotypes are not identical, as they are dynamic and transitory in the latter case.

**FIG 6.**
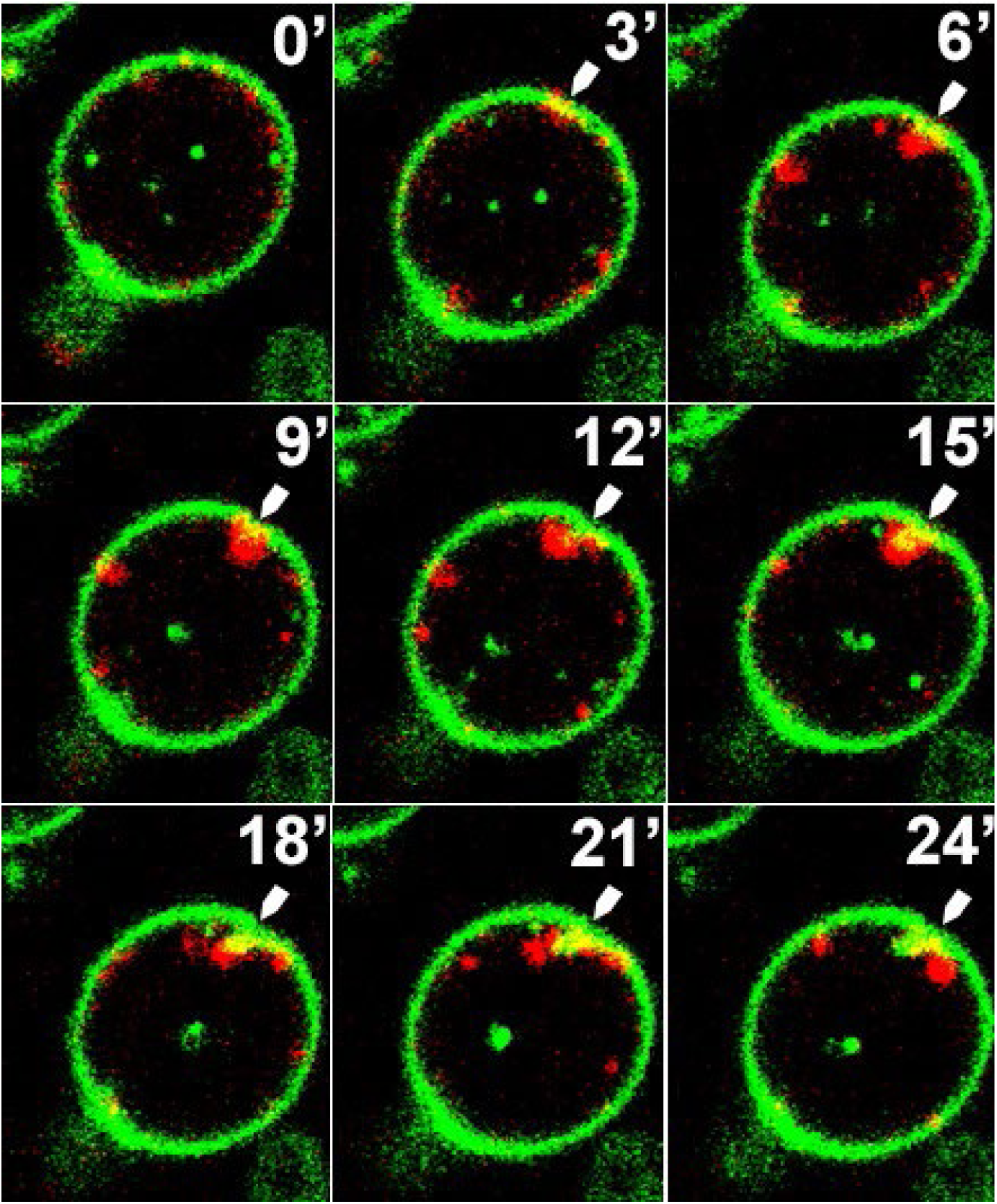
Exposure to the antifungal drug caspofungin mislocalizes actin along with PI(4,5)P_2_. Confocal live microscopy of wild-type cells co-expressing CaPHx2-CaGFP and ACT1-CaRFP showing a 3-minute time lapse images after treatment with 4xMIC of caspofungin. The white arrow points to a PM location where Act1p is recruited at 3’ and continues to increase till 12’, after which the PM with its PI(4,5)P_2_ starts to be pulled inside the cell to create an invagination. At 21’ and 24’, Act1p seems to dissociate from the PI(4,5)P_2_ invagination (See Supplemental Video 6).

**FIG 7.**
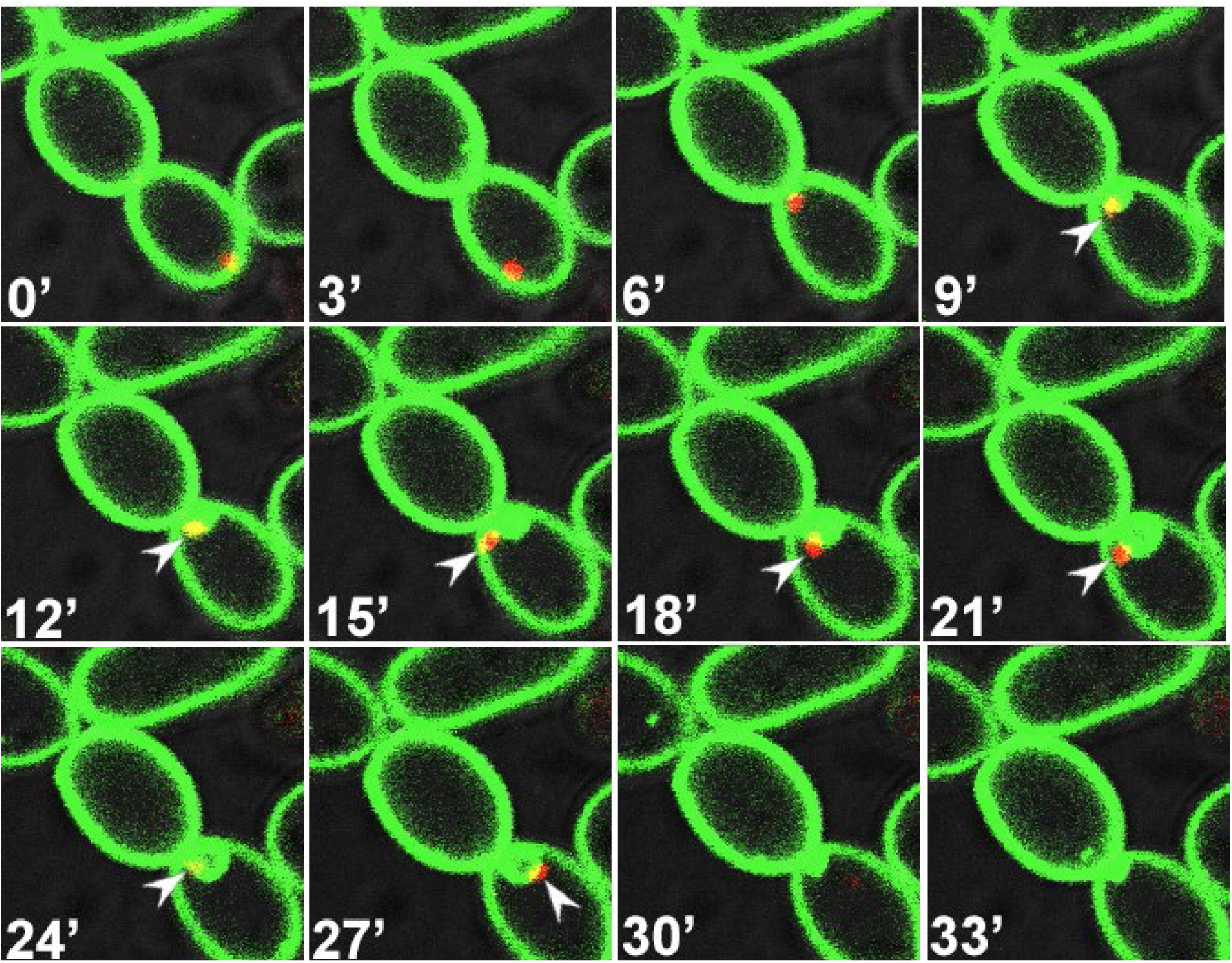
Exposure to the antifungal drug caspofungin mislocalizes Myo1 along with PI(4,5)P_2_. Confocal live microscopy of wild-type cells co-expressing CaPHx2-CaGFP and MYO1-CaRFP showing a 3-minute time lapse images after treatment with 4xMIC of caspofungin. The white arrowhead points to an instance of Myo1p mislocalization along with PI(4,5)P_2_, which happens right after the completion of cytokinesis (See Supplemental Video 7).

**FIG 8.**
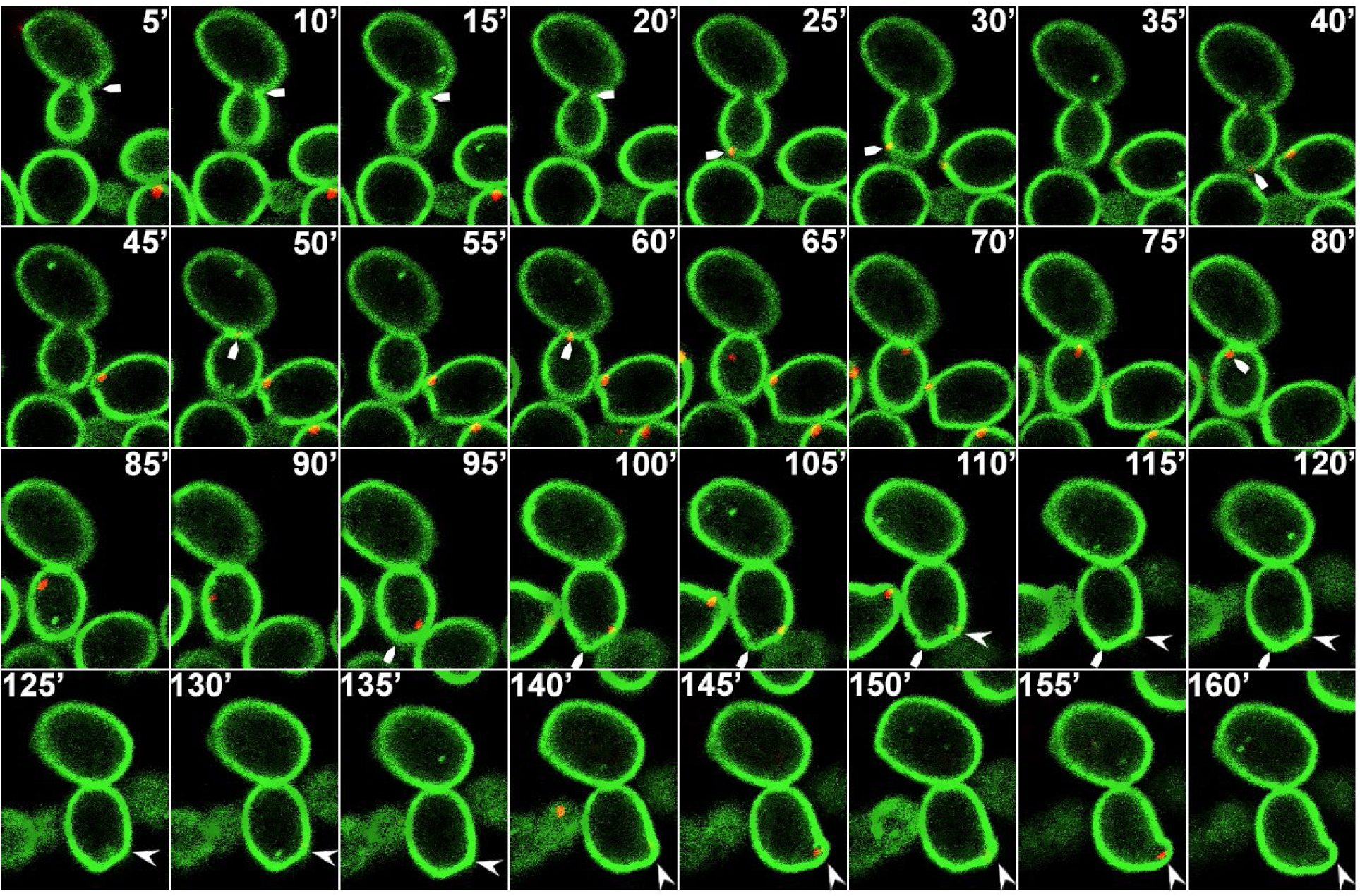
Exposure to the antifungal drug caspofungin mislocalizes Myo1p and causes other cellular defects. Confocal live microscopy of wild-type cells co-expressing CaPHx2-CaGFP and MYO1-CaRFP showing 5-minute time lapse images after treatment with 4xMIC of caspofungin. The white arrow indicates Myo1p localization being normal during steps of this cell division from 5’ to 80’ (indicated with white arrow), while PI(4,5)P_2_ shows multiple dynamic mislocalizations. During the following cell division, at 95’ the daughter cell from previous cell division starts a bud at a site indicated by white arrow, while Myo1 is localized to a different site where another bud is started at 110’ (white arrowhead). The initial bud seems to stall, but the second one continues the cell division with intermittent PI(4,5)P_2_ mislocalizations (See Supplemental Video 8). Note the unusually wide mother bud-neck.

**FIG 9.**
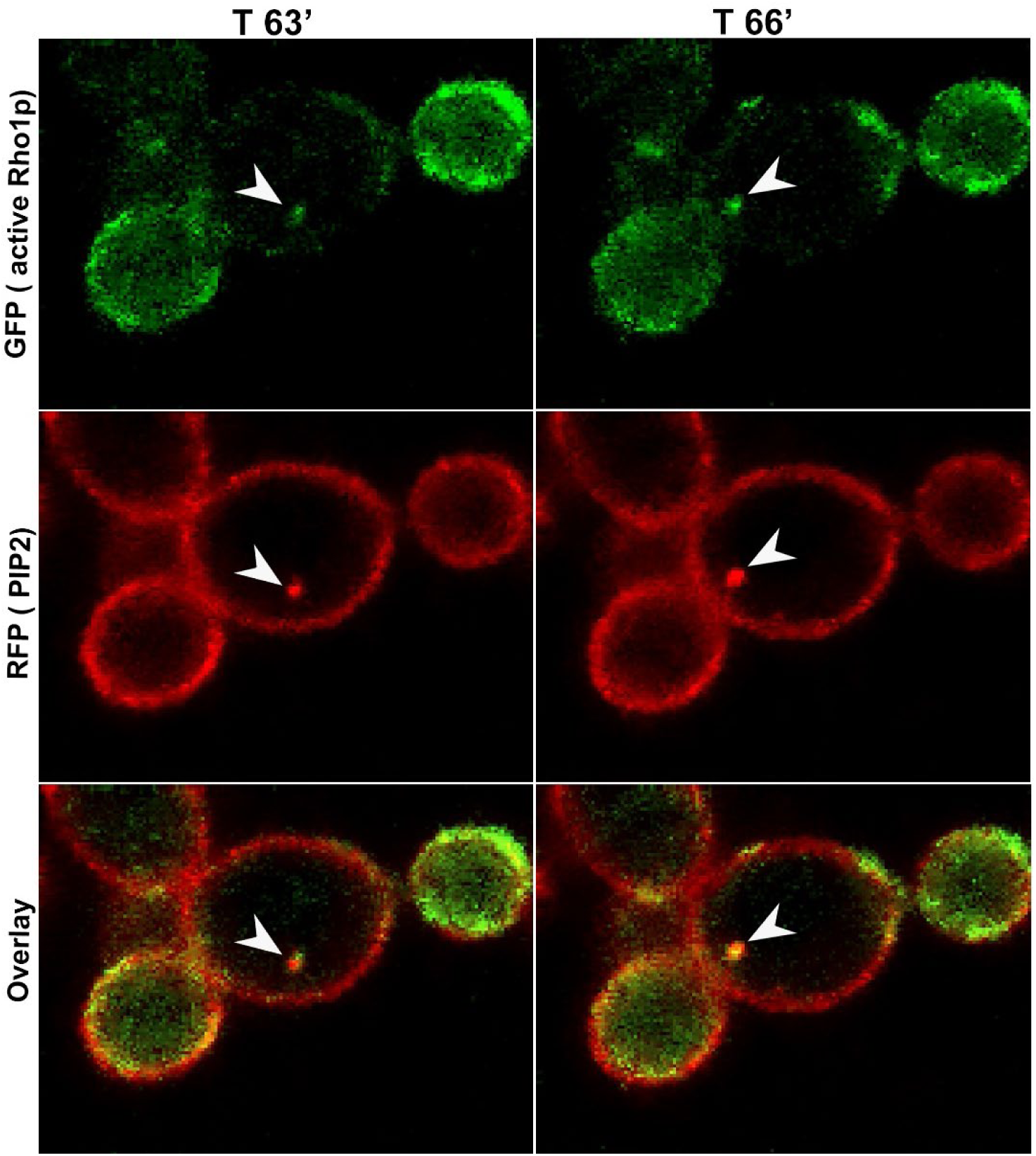
Exposure to Caspofungin mislocalizes active Rho1p along with PI(4,5)P_2_. Confocal live microscopy of wild-type cells co-expressing CaPHx2-CaRFP and PKC-RBD-CaGFP showing a 3-minute time lapse images at T 63 minutes and 66 minutes after treatment with 4xMIC of caspofungin. A dynamic mislocalization of active Rho1p together with PI(4,5)P_2_ is depicted in two different subcellular locations during a 6-minute period, indicated by white arrowhead.

## DISCUSSION

In previous work, we showed that exposure of *C. albicans* to caspofungin led to increased PI(4,5)P_2_ levels and septum-like invaginations of the PM that contained PI(4,5)P_2_, septins, and cell wall and PM constituents. These subcellular phenotypes were accompanied with dose-response caspofungin fungicidal activity and paradoxical growth at concentrations up to 4x MIC and > 4x MIC, respectively, and with levels of PKC-Mkc1 cell wall integrity pathway activation proportional to the degree of growth inhibition by the drug (8). The appearance of septation-like invaginations and presence of broad-based *C. albicans* mother-daughter bud necks suggested that caspofungin exposure might be associated with disordered cytokinesis. In the present study, we link PI(4,5)P_2_ regulation more directly to cytokinesis, and demonstrate that dysregulated PI(4,5)P_2_ and disordered cytokinesis are part of the natural response of *C. albicans* to caspofungin. Using live cell imaging, we show that PM patches and invaginations that concentrate PI(4,5)P_2_ are correlated in time and space to sites of cytokinesis in *C. albicans inp51* and *irs4* mutant cells in which PI(4,5)P_2_ 5-phosphatase activity is greatly attenuated. We then co-localize, in these mutant cells, PI(4,5)P_2_ with Act1p, Myo1p and activated Rho GTPase, the key components of the cytokinesis machinery. Finally, we demonstrate that cytokinesis is impaired in wild-type *C. albicans* SC5314 exposed to caspofungin, and that PI(4,5)P2 patches co-localizes with same cytokinesis components. Our data here and in previous studies support a model in which balanced PI(4,5)P_2_ regulation helps govern cytokinesis and cell wall integrity through a network that also includes septins and septin-regulating protein kinase Gin4, actin and myosin, Rho1 and the PKC-Mkc1 pathway (Fig. 10). Dysregulation of PI(4,5)P_2_ and this network, as seen with fungicidal caspofungin exposure and in *inp51* or irs4 mutants, is associated with disrupted cytokinesis and reduced cell wall integrity. PI(4,5)P_2_, the most abundant among the seven phosphoinositides in eukaryotes, is localized to the PM, where it interacts with effector proteins to perform diverse cellular functions, including ion channel regulation, actin cytoskeleton remodeling, endocytosis, exocytosis and phagocytosis. In addition, products of PI(4,5)P_2_ metabolism; conversion into PIP3 or degradation into inositol (1,4,5)-trisphosphate (IP3) and diacylglycerol (DAG), also relay further functions (4, 11).

**FIG 10.**
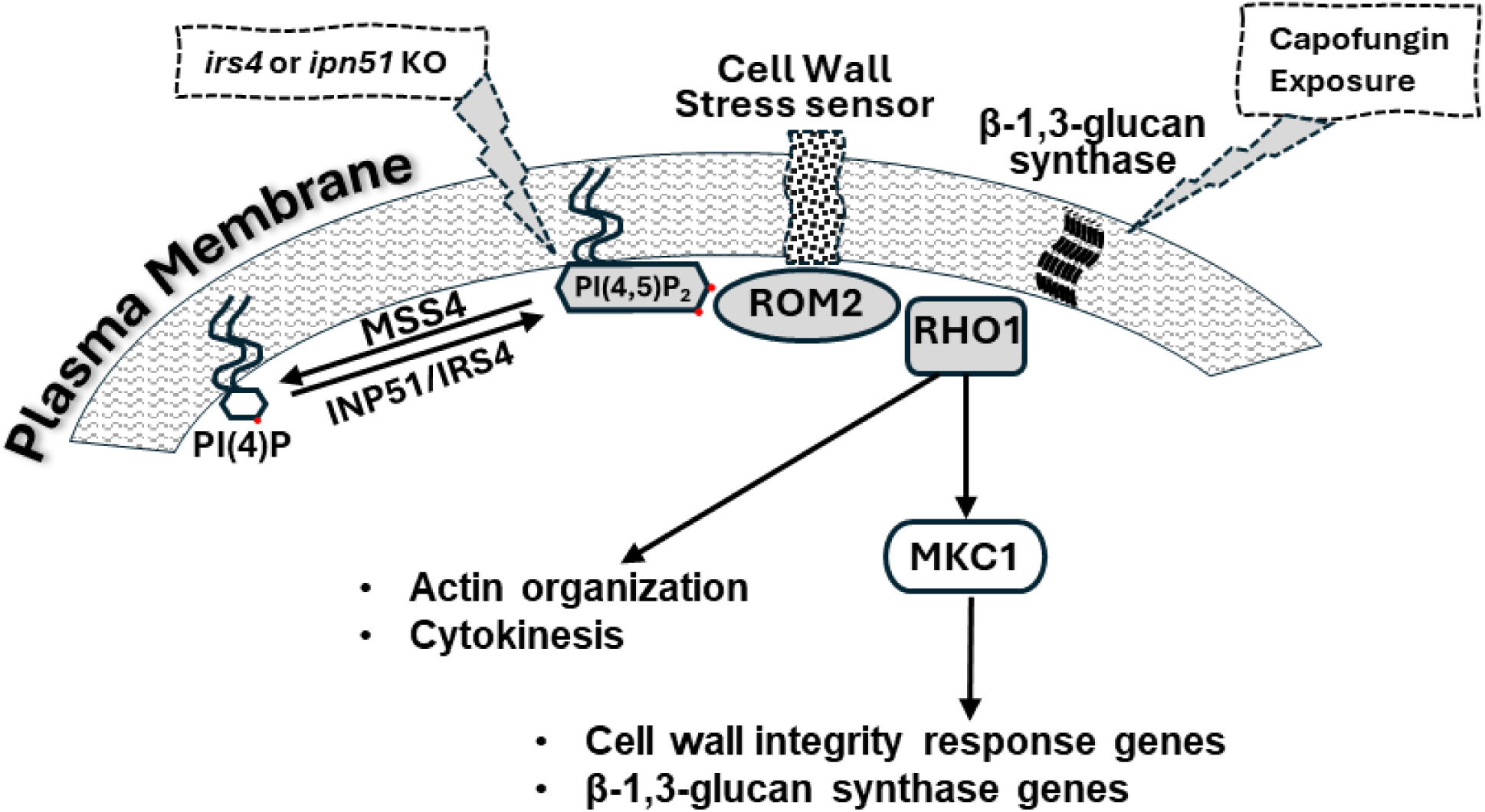
A schematic summary of the interplay between PI(4,5)P_2_ and Rho1p. At the PM, PI(4,5)P_2_ is synthesized from or degraded into PI(4)P by MSS4 or INP51, respectively. PI(4,5)P_2_ is able to interact with Rho1p either directly (Yoshida) or indirectly via ROM2 (Kobayashi). Interfering with this interplay between PI(4,5)P_2_ and Rho1p by either *IRS4* or *INP51* gene deletion, or by caspofungin exposure, leads to aberrant cytokinesis, perturbation in the cell-wall integrity pathway, and mislocalization of, not only PI(4,5)P_2_ and Rho1p, but also Act1p, septins, and Myo1p.

Phosphoinositides could be thought of as phospholipidic anchors that are implicated in virtually any function that involves cellular membrane dynamics. The type of membrane to which each phosphoinositide localizes dictates the type of its functions. It is then no surprise that PI(4,5)P_2_ is an essential molecule for cell life, and so is the gene encoding the protein which synthesizes PI(4,5)P_2_ from PI4P. This enzyme, which is a phosphatidylinositol 4-phosphate 5-kinase, was shown to be important for or regulate cytokinesis in *Schizosaccharomyces pombe* (its3 gene) (25, 26), and *Drosophila melanogaster* (fwd gene) (27). Several additional lines of evidence implicated PI(4,5)P2 in cytokinesis in various eukaryotes other than *C. albicans*. Overexpression of *MSS4*, an orthologue of its3, and Rho1 activation promote cytokinesis in budding yeast *Saccharomyces cerevisiae* even in the absence of an actomyosin contractile ring (28). Under normal circumstances, *S. cerevisiae* PI(4,5)P_2_ targets Rho1 to the site of cytokinesis, where the latter plays a central role in actomyosin ring assembly (28). Likewise, PLC activity is required for cytokinesis in flies spermatocytes from the order *Diptera* (29, 30). In mammalian cell lines, PI(4,5)P_2_ is enriched at the cleavage furrow and functions in adhesion of the PM to the contractile ring (31). Sequestering PI(4,5)P_2_ or disturbing its level in eukaryotes perturbs cytokinesis (for a review see (12)). This is the first study to explore associations between PI(4,5)P_2_, Rho1p, actomyosin ring and cytokinesis in *C. albicans*.

We previously showed that PI(4,5)P_2_ patches colocalized with septins; Sep7p and Cdc10p (8). The role of septins (Sep7p, Cdc10p, etc.), actin (Act1p), and myosin-II (Myo1p) in cytokinesis is well established (32). It has been shown that PI(4,5)P_2_ can directly bind the N-terminus of septins to promote their filament organization and stability (33, 34), and can affect actin polymerization and modulates actin cross-linking and regulatory proteins (35). Indeed, concentration of PI(4,5)P_2_ at the division site can facilitate anchoring of structural components of the actomyosin ring to the PM (35). Work by Yoshida et al. suggests PI(4,5)P_2_ could interact with a c-terminal polybasic sequence of Rho1p as part of a GEF-independent mechanism for targeting Rho1p to the bud neck (28). Our data present evidence that PI(4,5)P_2_ not only is a key player during cytokinesis, but its homeostasis is important for the correct spatial-temporal execution of this essential event in the cell cycle.

The echinocandin class of drugs, to which caspofungin belongs, impairs synthesis of cell wall β-1,3-glucan (36), by inhibiting β-1,3-glucan synthase enzymes encoded by *FKS* genes (37). It is currently unclear how caspofungin elevates levels of PI(4,5)P_2_ and results in its mislocalization. Since the drug is known to target FKS genes, the most obvious links to PI(4,5)P_2_ are the Rho GTPase RHO1, which regulates FKS genes (38–41), or the RHO1 activator, the GDP/GTP exchange factor ROM2, which possesses a PH domain (data not shown) and was shown in baker’s yeast to interact with PI(4,5)P_2_ (42). However, there are no data on possible mechanisms. It could also be a secondary effect to cell wall stress imposed by caspofungin, or a direct effect of the drug on enzymes responsible for PI(4,5)P_2_ synthesis or degradation. Once PI(4,5)P_2_ is dysregulated, it is no surprise to see an effect on septins, Act1p, Myo1p and active Rho1p, and ultimately defects in cytokinesis. We previously showed that caspofungin exposure rapidly causes a steady, dose-dependent increase in PI(4,5)P_2_ levels that continued throughout the 3 hours of live cell experiment, and which correlates with levels of fungicidal activity and paradoxical growth (8).

PI(4,5)P_2_, septins, Act1p, Myo1p and active-Rho1p mislocalization in wild-type cells exposed to caspofungin resembles that of *inp51* and *irs4* mutant cells in the absence of caspofungin exposure. However, these defects are transitory and more dynamic in caspofungin-treated wild type cells.

Taken together, our results allude to the possibility that the PI(4,5)P_2_ 5-phosphatase encoded by *INP51* along with its interacting partner Irs4p are critical during mitotic exit when final cell division is taking place, and that abnormal PI(4,5)P_2_ patches are remnants of imperfect cytokinesis. They also suggest that PI(4,5)P_2_, cytokinesis and cell wall integrity pathway responses that are associated with deleterious outcomes of caspofungin exposure are overexuberant expressions of normally protective responses likely governed by the interplay between PI(4,5)P_2_ and Rho1p (Fig. 10).

